# Saccadic compression of time as a marker for Developmental Dyslexia

**DOI:** 10.1101/2024.04.03.587978

**Authors:** Nicola Domenici, Alessia Tonelli, Cristina Ponente, Monica Gori

## Abstract

About 10% of the world’s population is dyslexic, experiencing reading impairments unrelated to cognitive deterioration. Due to its impact, identifying the mechanisms subtending dyslexia is paramount. However, while most research focused on the eye movements’ phenomenology, none investigated their perceptual, transient consequences. In fact, it has been shown that rapid eye movements (i.e., saccades) are accompanied by temporary distortions of space and time. Such distortions have been linked to the receptive fields’ predictive remapping, which anticipates the movement and compensates for the gaze’s displacement. Here, we demonstrate that dyslexic children show reduced flexibility in modulating temporal information around the saccadic onset. Moreover, accuracy oscillations within the delta band, phase-locked to the saccade’s onset, preceded transient temporal compression in typical readers. Conversely, no oscillatory behavior was observed in dyslexic participants, suggesting that the absence of transient temporal distortions originated from the mismatch between the anticipatory remapping and the saccadic onset.

## Introduction

As the reader approaches this first section of the manuscript, they surely have the impression that their eyes are smoothly gliding along each line of the text. Oppositely to such erroneous (but unavoidable) perception, while reading, we perform rapid, ballistic eye movements called saccades^1^, implying that our gaze “jumps” from one point to the other of the page. Because of these jumps, the retinal image of the text is constantly shifting as well, but these shifts are not matched with any perceived movement in the outside world (since the text is, hopefully, immobile as we keep reading). While this is still one of the most intriguing debates in vision science, a substantial amount of evidence suggests that such a degree of visual stability can be maintained via the predictive anticipatory remapping of receptive fields^2,3^: shortly before the saccade onset, receptive fields in the superior colliculus^4^, extrastriate cortex^5^, Frontal Eye Field (FEF) and Lateral Intra-Parietal (LIP) Areas^6,7^ relocate in the direction of the saccade, to pre-emptively compensate for the imminent movement. Interestingly, some cortical areas involved in this remapping mechanism – namely, the LIP – encode information about a stimulus’ timing and position^8,9^. It should not surprise the reader, then, that saccades are also accompanied by transient distortions of both spatial^10^ and temporal information^11,12^. Specifically, stimuli briefly displayed around the saccade’s onset are spatially mis-localized as being closer to the saccadic target and temporally underestimated^13^.

Since saccades are a central component of reading, they have been extensively studied in those who failed to acquire proper reading skills, i.e., dyslexic individuals. Indeed, many studies in past years highlighted atypical oculomotor behavior in dyslexic people, which show poorer binocular coordination^14–16^, lower saccadic accuracy^17^, and a smaller centre-of-gravity effect^18^ compared to typical reading peers. These crucial phenomenological manifestations have been linked to the magnocellular component of the visual system^19–21^ which specializes in perceiving rapid, transient information and is best suited to temporally encode external events^22–25^, being thus of great interest for the current work.

The magnocellular theory of dyslexia provides an insightful link between such a heterogeneous mosaic of evidence^26–28^, which is focused on the perceptual, rather than phonological, deficits associated with it^29^. The infrastructure of this exhaustive theory was built over a significant amount of studies turning the spotlight on functional^30^, genetic^31^, and anatomical^32–35^ differences involving the magnocellular system in dyslexic individuals. A key point of this theory, at least for our study, is the neuroanatomical predisposition of the magnocellular stream in encoding rapidly-changing, transient information, suggesting that its defective development should determine a significant impairment in temporal perception. Unsurprisingly, this has been found in people with dyslexia for more than 40 years^36–40^.

In light of the implications of the magnocellular theory of dyslexia, here we tried to expand its boundaries focusing on the saccades’ perceptual consequences rather than just investigating their mechanical components. We speculate that, since magnocellular processing is impaired in dyslexia, saccadic compressions should reflect this impairment by highlighting reduced efficiency in modulating visual time shortly before an eye movement occurs. Moreover, this inefficiency should leave a mark considering how perception, in the typical adult brain, fluctuates before the saccade. In fact, many studies demonstrated that perception oscillates shortly before the onset of a saccade and that these oscillations are phase-locked to the start of a movement^41–45^. Considering that neural oscillations play a key role in synchronizing action and perception since the early phases of motor planning, they should be as crucial in timing the predictive remapping of the receptive fields along the visual pathway. In addition to that, dyslexia has often been associated with atypical eye movements^46^, and one of the primary brain areas tuned for eye-movement parameters is the LIP^47,48^ – in which the predictive remapping of the receptive fields occur the most^49^.

To combine all these different frameworks, we thus measured the saccadic compression of temporal information in both dyslexic and typical-reading children, developing a gamified version of a series of paradigms already used in adult participants^10,12^. Considering our premises, we speculate that dyslexic children will show less flexibility in modulating time perception around the start of the saccade when compared to typical readers of the same age. Moreover, investigating fluctuations in timing perception before the saccadic onset will shed light on whether there is any difference between groups in how the brain prepares prior to a saccade. If no perceptual oscillations will anticipate saccades in dyslexic children, it is feasible to assume that a missed synchronization between visual perception and the execution of the motor command occurred, thus resulting in a mismatch between the saccade itself and the remapping of the corresponding receptive fields.

## Results

### Saccadic compression of time

In the current study, we asked children to perform saccadic movements subtending 20° of visual angle, from an initial fixation point placed 10° leftward to the observer to a saccadic target placed 10° on their right. A first test interval, delimited by two bars briefly displayed around the initial fixation point, was presented either before or around the saccadic onset. After participants performed the saccade, a second (reference) interval was displayed, in a similar fashion, around the saccadic target. In order to complete the task, participants had to report which one of the two intervals lasted longer (Figure 1).

**Figure 1:**
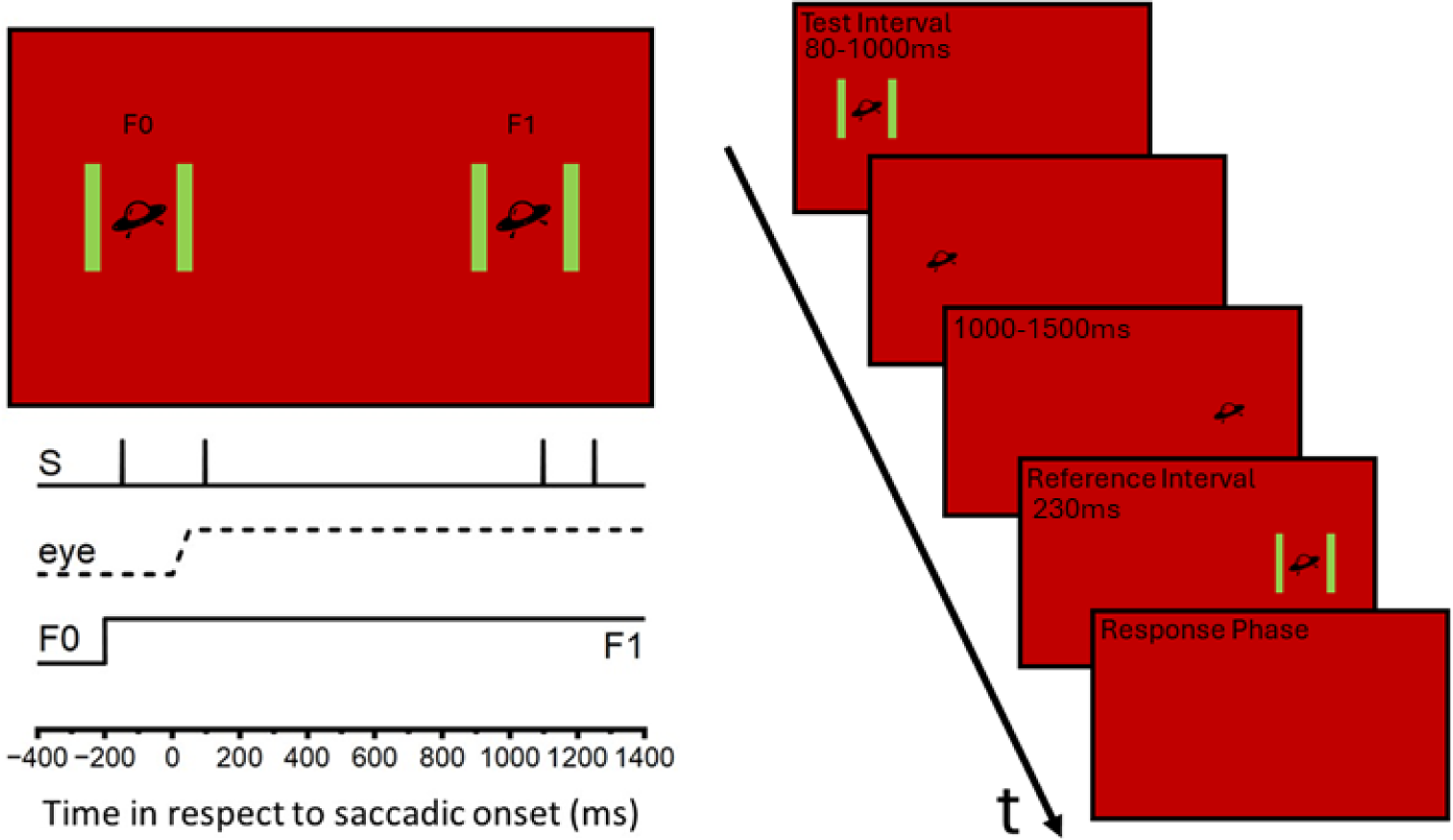
Experimental design, including the representation of visual stimuli (S) and their timing (right panel). During each trial, an empty Test interval, delimited by two sequentially displayed green, vertical bars (2 × 10° of visual angle), was displayed on a red background around the fixation point (F0). Bars were separated by an interval ranging from 80 to 900ms and displayed for one frame (7ms) before or around the saccadic onset. As soon as participants had maintained fixation on F0 for one second, the fixation point disappeared, and a fixation target (F1) simultaneously appeared rightward placed at 20°. After participants executed a saccade towards F1, a 230ms reference interval was displayed around the target. To conclude the trial, participants were asked to indicate via keypress which interval lasted longer. On the right panel is depicted the sequence of events that characterized a sample trial, in which the test interval was displayed before inducing a saccadic movement. After fixation was maintained on the saccadic target for 1000-1500ms, the Reference interval was displayed. Finally, participants provided an unspeeded response via keypress to indicate which interval lasted longer, so that the next trial could start.

Considering this work’s premises, to scan for different perceptual patterns between dyslexic and typical-reading children, we directly evaluated how performance changed in the two groups before the saccade’s onset. Intriguingly, it has already been shown that perceptual oscillations within the delta band (0.5-4 Hz) systematically occur before executing a voluntary eye movement^42,43^, suggesting that vision is rhythmically modulated before a saccade. Specifically, Di Benedetto and Morrone demonstrated that saccadic suppression^50,51^ – the abrupt sensitivity reduction that characterizes saccades – is embedded in phase with these oscillations^42^. Therefore, we decided to investigate how performance in the temporal task developed up to 1000ms before the saccade’s onset, looking at how – and if – temporal sensitivity (resembled by the proportion of correct responses) changed according to a well-defined pattern. Due to the low number of trials per participant, we investigated oscillatory behaviour on the aggregate data, combining performance across all participants within each group. We then binned all trials (using non-overlapping 100ms bins) and obtained 10 points to find the best-fitting sinusoidal waveform (Figure 2A).

**Figure 2:**
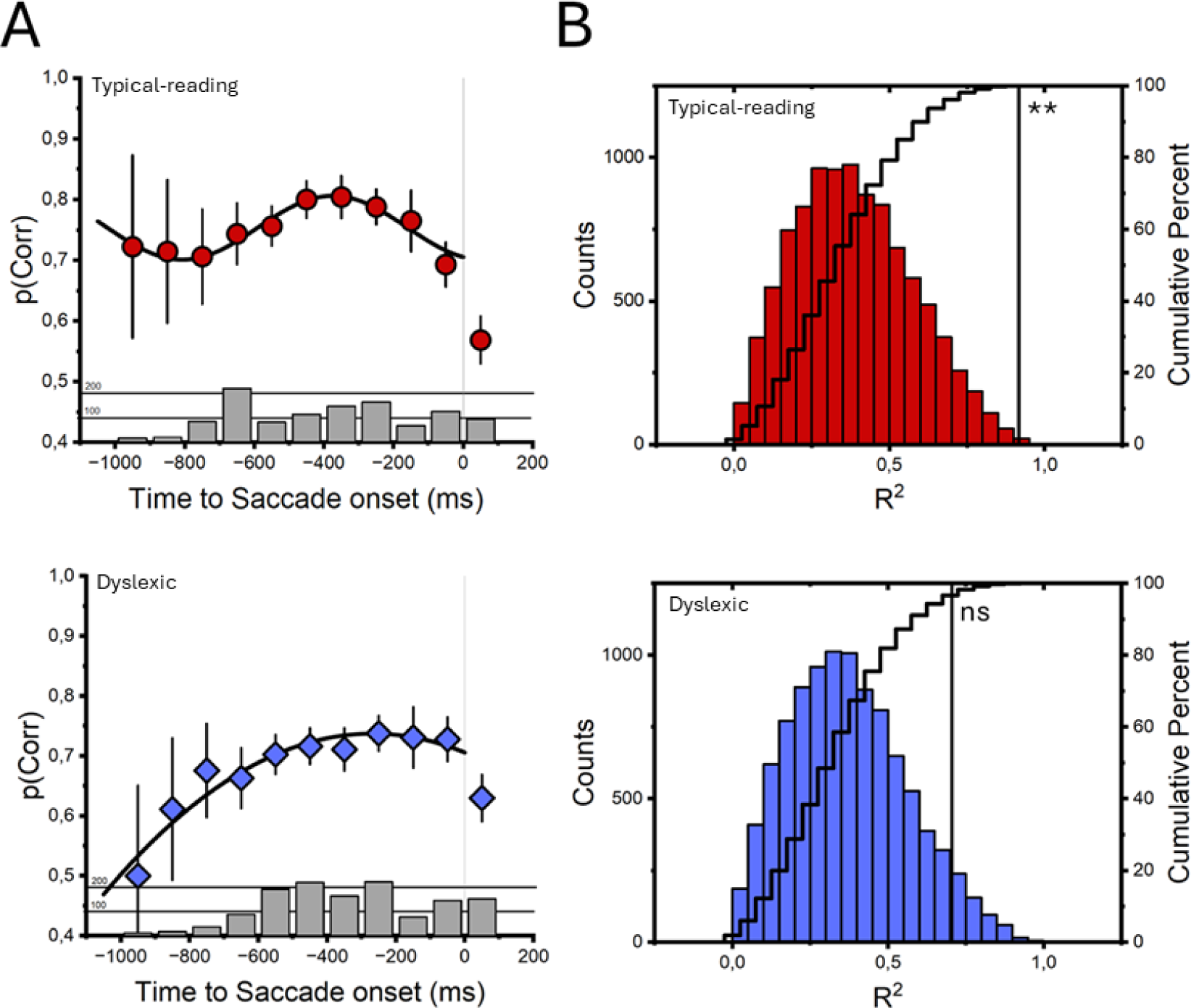
A) Oscillation of temporal sensitivity up to 1000ms before the saccade’s onset considering aggregate data of typical-reading (top panel, red scatter points) and dyslexic (bottom panel, blue scatter points) participants. Error bars indicate ± SEM for any given bin, while the black line represents the best sinusoidal fit obtained considering all 10 bins. Bar plots report the number of trials per bin, while grey, vertical lines identify the saccadic onset. B) Distributions of R^2^ obtained by fitting sinusoidal functions on surrogate datasets for both typical-reading (top panel, red bars) and dyslexic (bottom panel, blue bars) children. To evaluate if the empirical sinusoidal fit was statistically better than a fit obtained stochastically, we compared its R^2^ (pinpointed by the vertical lines) against the ones obtained by fitting randomly generated surrogate datasets. Stairs report the cumulative percent of the R^2^ frequency distribution (** = p-value<0.01).

We thereby evaluate the best sinusoidal fit approximating the data, which yielded extremely reliable goodness-of-fit parameters (R^2^=0.917, p=0.0012; see the *Fitting Procedures* section for additional information).

To assess the model’s ability to truly represent the data, we repeated the analysis on 10,000 surrogate datasets obtained by randomly shuffling, for each trial, the temporal distance between the stimulus presentation and the saccadic onset (to prevent spurious effects caused by the trials’ temporal distribution, as well as to maintain the same number of trials per bin). For each permutation, we fitted the best sinusoidal function (keeping phase, amplitude, and frequency as free parameters to achieve correction for multiple comparisons) on the same 10 bins. Then, we compared the empirical *R*^2^ (obtained through the sinusoidal fit on the empirical data) with the distribution of *R*^2^ obtained with the surrogate datasets (Figure 2B). Statistical significance was evaluated considering the probability that the empirical *R*^2^ distribution was higher than any of the surrogated *R*^2^ (with α=0.05).

The results of our fitting procedure on typical-reading participants’ aggregate data showed not only that temporal sensitivity significantly fluctuated before the saccade’s onset (Figure 2A, upper panel) but also that our sinusoidal model well-captured such oscillation (Figure 2B, upper panel), being significantly more reliable than the permuted fit performed on surrogate data (p=0.0013). Interestingly, the best-fitting sinusoidal function was phase-locked with the saccadic onset and showed a frequency of about 1.16 Hz (which fell within the delta range), confirming the results of previous studies investigating saccadic suppression in adults^42,43^.

To evaluate whether delta oscillations characterized pre-saccadic temporal sensitivity in Dyslexic, we implemented a similar modeling procedure. Notably, the best sinusoidal function fitted in the 10 bin points failed to converge and, although performed significantly better than a constant line (R^2^=0.706, p=0.002), its shape already suggested that comparable oscillatory behavior failed to characterize dyslexic participants (Figure 2A, lower panel). Regardless, delving further into the analysis pipeline, we found that the R^2^ obtained with the empirical data was not significantly higher than the ones obtained through random generation of surrogated datasets (p=0.053), further suggesting that no oscillatory behavior characterizes temporal performance before the saccadic onset in Dyslexic (Figure 2B, lower panel).

These results were expanded by fitting an ensemble of psychometric functions into the aggregated data,considering in separate analyses all trials characterizing one specific bin at a time, and obtaining a psychometric curve for each time bin (Figure 3). Within the current study, psychometric curves indicate the probability of indicating the Test interval as the longer ones, expressed as a function of the difference between the Test and the Reference. By following such statistical procedures, we were able to investigate how performance changed both before and after the saccadic onset just comparing the inflection point of each curve (indicating the Point of Subjective Equality, PSE). Notably, in typical-reading participants the curve with the highest PSE was the one obtained when fitting data in the 50ms bin, suggesting that the highest temporal compression was achieved shortly after the onset of the saccade. Conversely, the 50ms psychometric curve obtained with Dyslexic participant showed a much smaller PSE, indicating that the temporal compression induced by the saccade was significantly limited.

**Figure 3:**
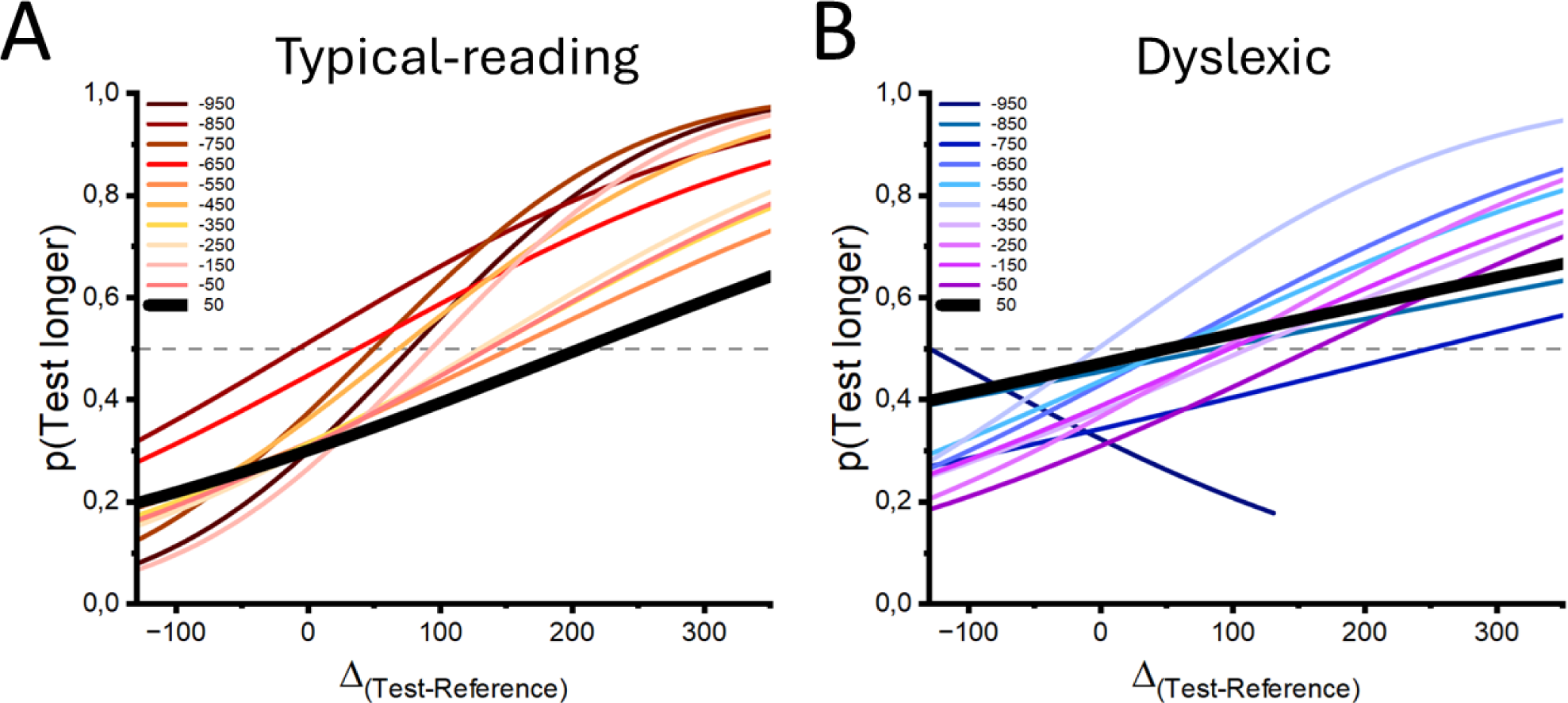
A) Ensemble of psychometric curves obtained by fitting a psychometric function into all trials for each bin, considering typical-reading participants. Here, psychometric curves report the probability of indicating the Test as the longer interval, expressed as a function of the temporal difference between the Test and the Reference. The black psychometric curve represents the performance around the saccadic onset (−50ms), where the temporal compression is expected to peak. In typical-reading children, the highest PSE (visualized as the inflection point, corresponding to the temporal difference between intervals for which performance is at chance level) among all bins was observed when considering trials immediately before the saccadic movement, suggesting that transient distortions of temporal information characterized performance around this timeframe. B) Similar analysis performed on Dyslexic children. Note how the psychometric curve obtained with all trials making up the −50ms bin is not associated with the highest PSE, indicating that dyslexic children were less flexible in modulating temporal information around the saccadic onset

Finally, to provide additional statistical foundation to our results, we run a Bayesian Logistic Regression model at the single trial level, including as dependent variable the correctness of the response (1 = correct, 0 = incorrect), and as a factor both the Group (Typical reading vs. Dyslexic children) and bin (−950 vs. −850 vs. - 750 vs. −650 vs. −550 vs. −450 vs. −350 vs. −250 vs. −150 vs. −50 vs. 50). We included in the analysis both the Age (expressed in months) and the reading speed as covariates, and add them to the null model in order to control for their influence. Overall, our analysis highlighted that the best model describing the data included both factors (Bin + Group), showing a BF_10_ of 6.183 when compared to the null model (suggesting that our data were six times more likely to occur under the alternative than the null hypothesis). Posterior odds analyses showed inclusion BFs of 3.98 and 6.36 for the Group and Bin factor, respectively, suggesting a relevant increase for the model that included both predictors. Taken together, these results concreted all findings discussed until now, suggesting that not only there were differences in performance between groups, but also depending on the temporal difference between the stimulus presentation and the saccadic onset. Posterior summaries of coefficients are reported in the *Posterior odds for the logistic regression* section.

### Saccadic compression of space

To investigate whether the reduced flexibility in modulating information around the saccadic onset found in dyslexic participants was domain-specific, we ran a second experiment aiming at inducing spatial transient distortions within the same timeframe. To minimize biases due to familiarization with the setup and design, the order of presentation of the current task and the one described in the previous section was counterbalanced across participants.

In this second task (Figure 4), a green vertical bar (i.e., the target) was briefly presented either before, during, or after the saccade’s onset, placed in one of three possible positions (−10°, 0°, or 15° in respect to the participant’s head position). After the execution of the saccadic movement, a green vertical bar, identical to the target, appeared on the screen. Participants were then asked to control it to pinpoint the position in which they believed the target was displayed.

**Figure 4:**
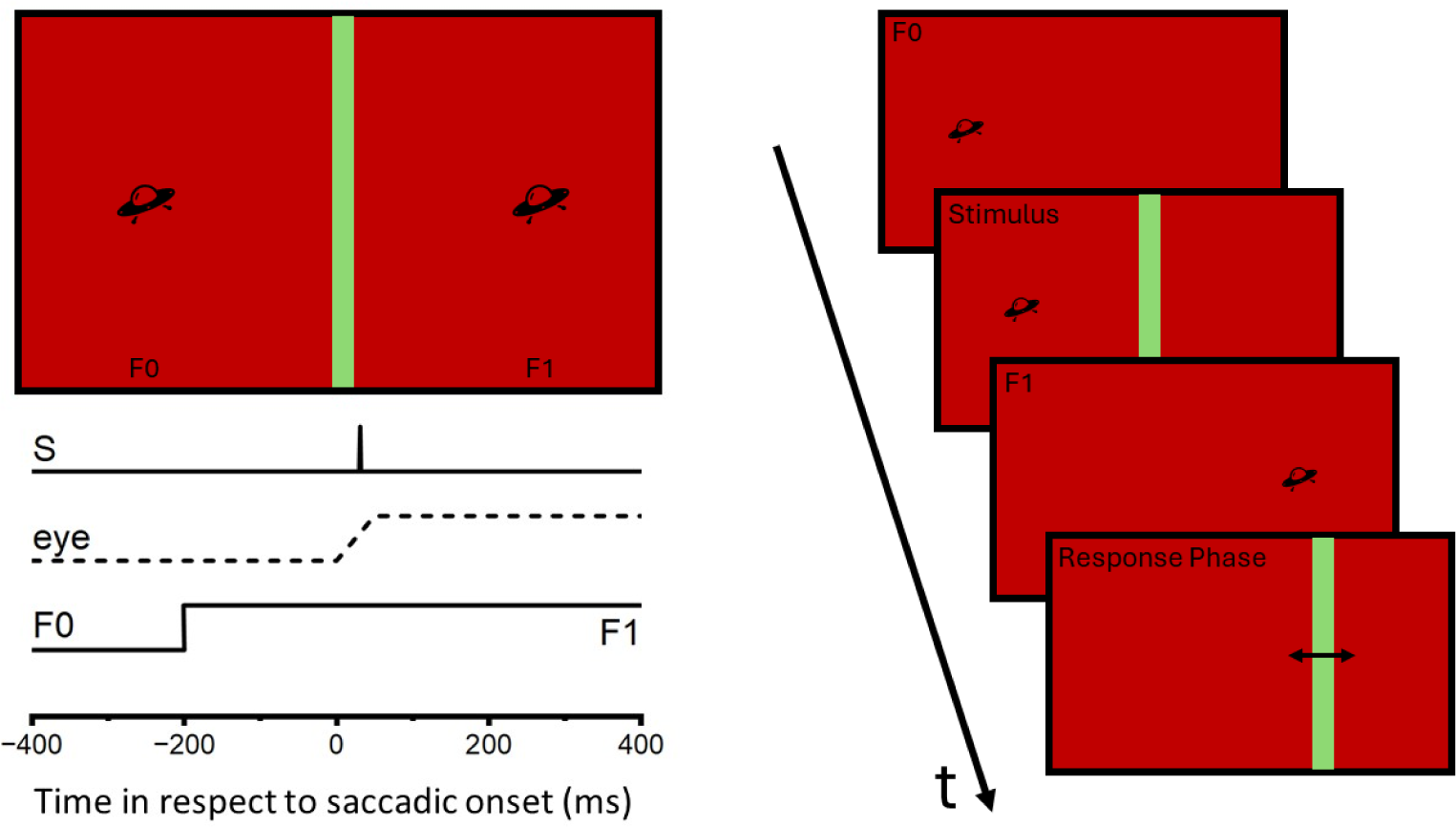
Experimental design for the spatial task, including the representation of visual stimuli (S) and their timing. During each trial, a single vertical bar (2 × 10° of visual angle), was displayed on a red background in one of three possible positions (−10, 0, and +15° of visual angles in respect to the observer). The bar was displayed for one frame (7ms) before, around, or after the saccadic onset. After participants maintained fixation on F0 for one second, the fixation point disappeared, and a fixation target (F1) simultaneously appeared placed 20° to the right. To conclude the trial, participants were asked to report the perceived position of the bar by controlling a similar one with the mouse (Response Phase in the right panel). The right panel reports the temporal evolution of a trial.

To investigate how saccades shaped spatial perception in dyslexic and typical-reading children, for each trial we evaluated the spatial mislocalization by calculating the difference in position (reported in ° of visual angle) between the participant’s response and the actual physical position of the bar. To maintain consistency across all three bar positions, we multiplied the difference by minus one when the bar was placed rightward to the saccadic target (at 15° of visual angle to the participant’s point of view). By doing that, we ensured that values greater than 0 always indicate mislocalization towards the saccadic target (i.e., attraction), while values lower than 0 indicate mislocalization away from the saccadic target (i.e., repulsion).

Akin to the analysis performed for the temporal task, we investigated to what extent perceptual oscillations anticipating the saccade characterized performance in the spatial task (n=51, 26 dyslexic and 25 typical-reading children; from the final analysis, we excluded trials in the −850ms bin due to the low number included in it: 3 in the typical-reading and 2 in the Dyslexic group). Differently from the temporal task, we failed to highlight any oscillatory behavior preceding the saccade (Figure 5), even though both typical-reading and dyslexic children showed significant saccadic spatial distortions around the saccadic onset. For typical-reading participants, the best-fitting sinusoidal function was not statistically significant (*R*^2^ = 0.44, p = 0.47), showing the following parameters: *y0* = −1.1682*, A* = 1.0838*, ω* = 127.3459, Φ = 137.4562. Similarly, no perceptual oscillations were observed in dyslexic children (*R*^2^ = 0.56, p = 0.3) before the start of the saccade (best sinusoidal function’s parameters: *y0* = −1.0975*, A* = 1.3062*, ω* = 137.8103, Φ = 164.8093).

**Figure 5:**
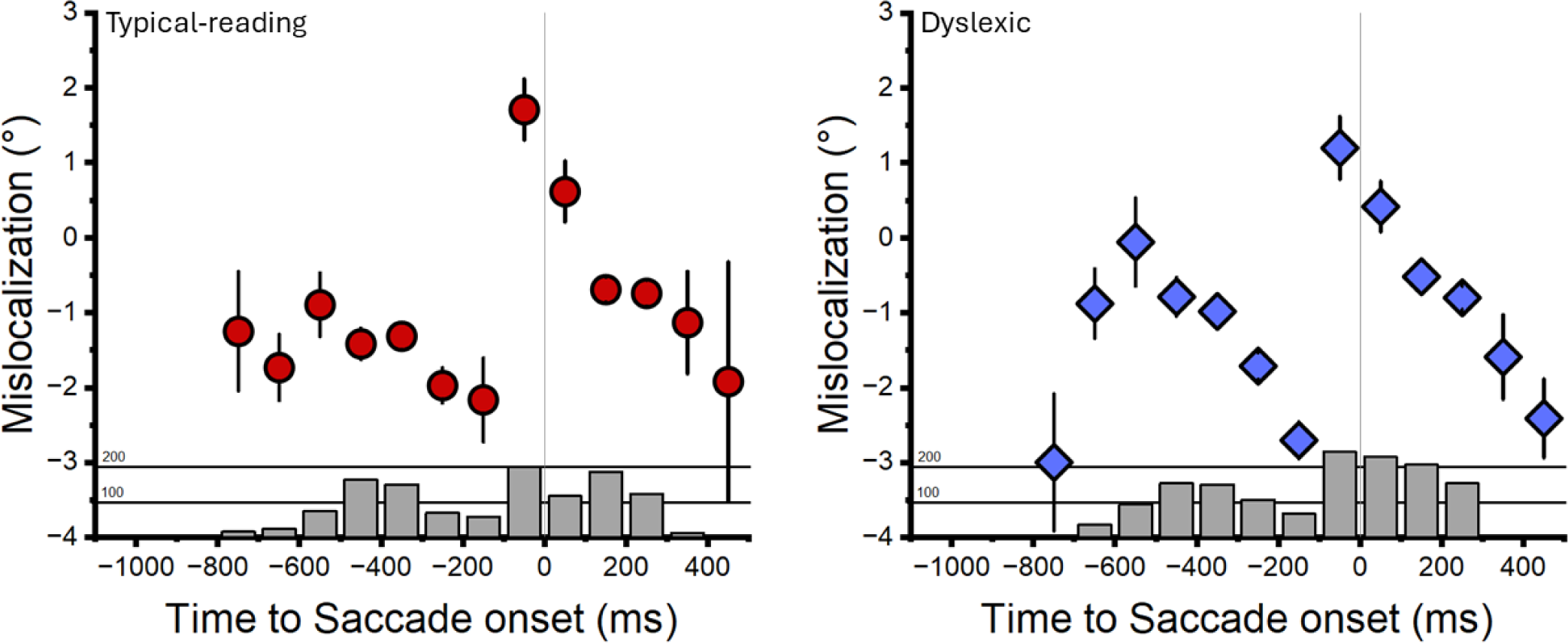
Perceptual oscillations for the spatial task. In none of the two groups the best sinusoidal fit achieved statistical significance, suggesting that before a saccade spatial and temporal information might be encoded differently. Nonetheless, considering the limitations concerning the design discussed in the section, further investigation on the matter is surely needed. In both graphs, error bars indicate ±SEM, while grey bars below each bin indicate the number of trials included in that bin.

Despite the lack of tangible oscillatory behavior prior to saccadic onset when modulating spatial information, we ran a Bayesian ANCOVA at the single trial level, considering as dependent variable the mislocalization and as factors both the Group and the Bin (similarly to what already done for the analysis of the temporal performance). We also added as covariates the Age and the Reading Speed, which were added to the null model in order to control for them. Overall, our analysis highlighted that the best model approximating the data was the one including only the Bin factor, which showed a BF_10_ of 2.023 × 10^26^ when compared to the null model. Interestingly, when compared to the best (only Bin) model, both the model including only the Group (BF_10_ = 1.435 × 10^-27^) and the one including the interaction between factors (BF_10_ = 0.001) showed BF_10_ significantly lower than 1/3, indicating crucial evidence that the Group factor was not determining changes in performance, nor that it was interacting with the factor Bin. As a follow-up analysis, we ran post-hoc comparisons on the factor Bin to test whether the spatial mislocalization around the saccadic onset was different from the ones experienced before the start of the eye movement. Our results further demonstrated that the mislocalization peaked around −50ms (Figure 6), as the spatial distortions found within this time window (1.43°) higher than all the mislocalizations observed before the saccadic onset (all BF_10_>3) but the one within the −750ms bin (mostly due to the response variability measured in the latter). All post-hoc comparisons are reported in Table T4.

**Figure 6:**
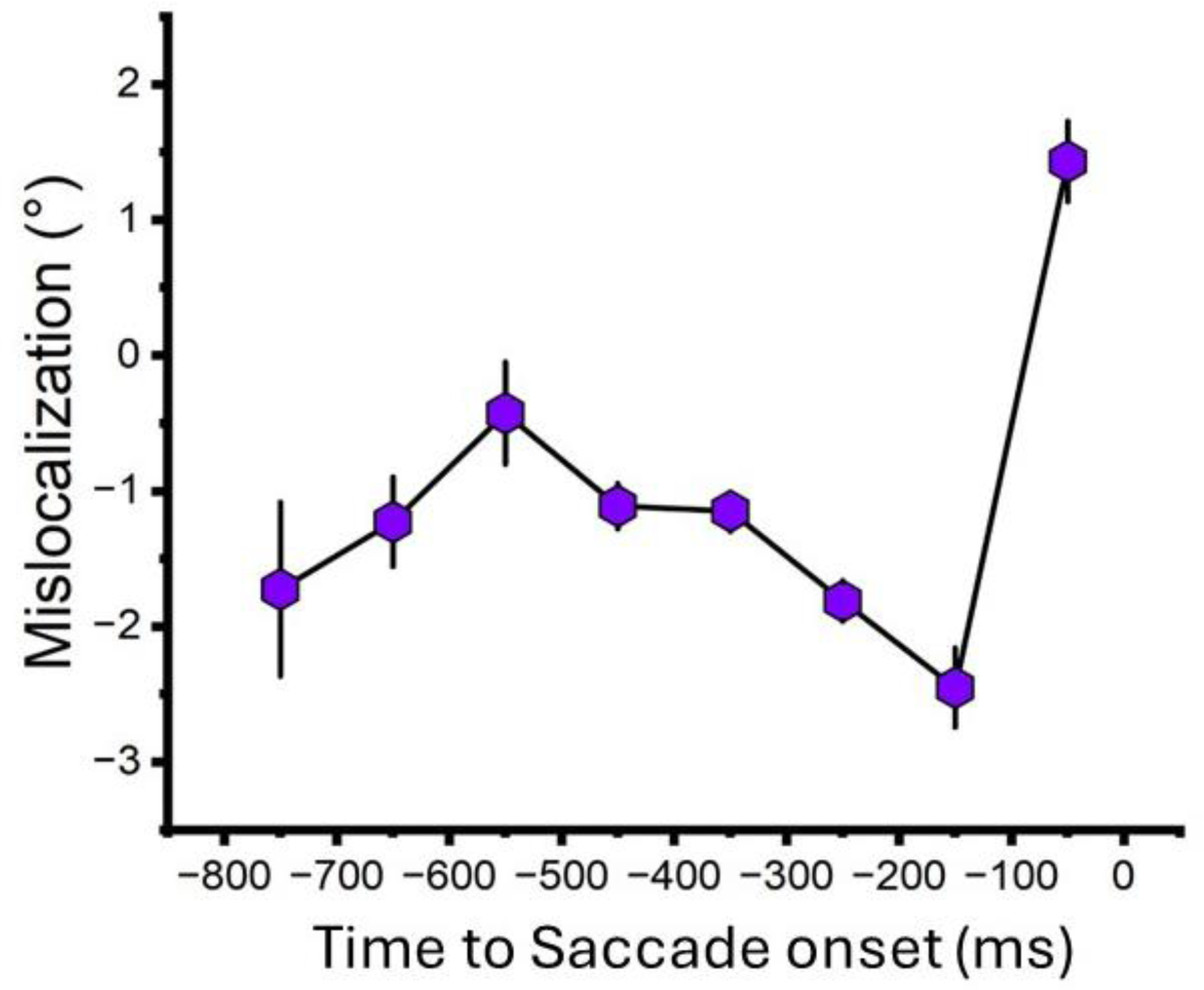
Mislocalization of the bar expressed as a function of the Time to saccadic onset, including all participants from both groups. Error bars indicate ±SEM.

## Discussion

With the current work, we investigated the ability to modulate temporal and spatial information shortly before and after a saccadic movement in children with and without developmental dyslexia. Our results highlighted relevant differences when testing children’s ability to evaluate the duration of an empty interval displayed around the start of the saccade, while no difference was found when considering spatial transient distortions. Interestingly, while typical-reading children experienced a significant compression of the interval’s perceived duration in the perisaccadic window, dyslexic participants showed lesser flexibility, reporting more veridical durations regardless of the saccade’s execution.

Although the origin of saccadic compressions is still an object of debate, there is strong consensus^12,51^ that these transient distortions result from the receptive fields’ predictive remapping occurring mainly (but not exclusively) within the dorsal stream^3–7^. Both saccadic suppression^50^ and compressions^12,52^ might function as a dynamic compensatory mechanism to maintain visual stability across eye movements. Interestingly enough, perceptual consequences of saccadic movements have been investigated not only between^12,52^ but also within participants^13,53^, showing that saccades affect both spatial and temporal processing in a similar fashion^13^ presumably due to the recruitment of the same neural correlates and the existence of visual receptive fields oriented in spacetime^54^.

In the current work, we found evidence supporting the existence of temporally sensitive receptive fields in typical-reading participants, while also proving that similar dynamics might fail to develop in dyslexic children (Figure 2A). Hence, we proceeded to observe how temporal sensitivity oscillated before the saccade’s onset, investigating fluctuation in performance suitable to explain such perceptual behavior. In fact, changes in both sensory accuracy and precision preceding a saccade have already been documented in the past^42,43^, showing how visual perception and saccadic preparation are tightly coupled to temporally match the saccade’s onset. Although a similar link has not been described yet for what concerns saccadic temporal compression, it is plausible to assume that – being a similar by-product of eye movements^54^ – the latter follows analogous principles due to the required temporal conjunction with the start of the saccade.

In line with this prediction, we found that, in typical-reading children, temporal sensitivity oscillated up to one second before the saccade’s onset, showing rhythmical fluctuations within the delta range (between 1 and 4 Hz, Figure 2A, upper panel). However, dyslexic children showed a much less predictable performance, as their temporal sensitivity did not oscillate before the saccadic onset (Figure 2A, lower panel). To date, there is still debate on the neural mechanisms behind such an oscillatory behavior. However, it has been suggested that pre-saccadic changes in perception are not only synchronized with the execution of a saccade, but also determined by the interaction between the endogenous, intrinsic brain oscillations and the exogenous source of stimulation^43^. Thus, given the number of saccades performed during natural reading, it is plausible to assume that the brain tries to constantly phase-lock inherent oscillations with the visual input, timing the formers at each eye movement.

Recently, Pan and colleagues have tightened the link between reading and brain oscillations, which showed that, when reading, saccades are phase-locked to alpha (∼8-13 Hz) but not to delta oscillations^55^. While these results seem to collide with our findings at first, oscillations within both ranges have been related to changes in either perceptual sensitivity and bias, considering the delta and the alpha range, respectively^43^. Moreover, while the execution of a saccade appears to be phase-locked to the alpha oscillations, its perceptual consequences might be phase-locked to the delta ones, further suggesting that people with dyslexia exhibit atypical neural patterns not only mechanically, when timing eye movements, but also physiologically, when modulating incoming visual information and preparing for the movement. This consideration naturally raises the following question: why did dyslexic children fail to transiently modulate visual temporal information within the perisaccadic window? Considering the neural mechanisms plausibly at the basis of saccadic compressions (namely, the predictive anticipatory remapping of the receptive fields), we argue that dyslexic failed in coordinating the receptive field’s shift occurring before the saccade and the saccade’s onset itself, resulting in a temporal misalignment which is crucial in determining the absence of transient distortions. Notably, our explanation is supported by the absence of oscillations in the perceptual behavior preceding the saccade found in dyslexics children, which suggests that the brain is somehow ‘less aware’ of when the saccadic movement will start. Unsurprisingly, various studies have already proved that oscillations in visual performance are synchronized with the onset of action^41,44,45,56^, indicating that action and perception are coupled since the early stages of motor planning.

On a final note, why should the absence of saccadic transient distortions observed in dyslexic children involve only temporal information? Considering the extensive literature on Dyslexia that thrived in the last 40 years, here we speculate that such a domain-specificity reflects the magnocellular impairment characterizing the clinical picture of this specific learning disability^20,26–29,32,33^. Given their fast response properties, magnocellular neurons are indeed well-suited to capture rapid changes in the environment^24^, and impairment at the magnocellular level is often linked to reduced temporal sensitivity – such as in dyslexic individuals^36–40^. Such considerations might thus explain, to some extent, why dyslexic children experience reduced flexibility in modulating temporal information around the saccadic onset. Additionally, magnocellular cells mainly project to the dorsal stream^57^, the pathway along which predictive remapping primarily occurs^4–7^, suggesting that the information conveyed by these cells was originally included within the shifting receptive fields. Therefore, the intrinsic noisier temporal processing typical of Dyslexia hampers perception fluctuations, making the action-perception synchronization far more difficult to set properly – even when significant differences in temporal abilities are not as evident at first. Consequently, the anticipatory shift of the receptive fields is temporally misaligned with the saccade, and visual timing maintains constancy across the saccade’s execution.

For what concern the spatial distortion induced by the saccadic movement, we would like to further expand its interpretation considering two major points. First and foremost, the response in the spatial task was collected directly by each young participant, who could move the mouse at will before registering the input. Such design, while more engaging, might have determined the occurrence in the response of a hard-to-quantify noise due to its motor component. Motor noise was normalized by comparing the mislocalization along all trials, indicating that the increase in mislocalization observed around the saccadic onset is not an epiphenomenon arisen from the mere participant’s inability to pinpoint the exact position of the target. Nonetheless, such noise might had a more relevant impact on our results when trying to highlight smaller effects, such as perceptual oscillations. Secondly, the lack of trials with a Time to saccadic onset lower than −800ms surely hindered the analysis itself, as it forced us to scan performance over a smaller time range and with a reduced number of bins.

We firmly believe that the study of perceptual oscillations subtending spatial saccadic distortions should be expanded in the future. However, one of the primary goals of this second experiment was to highlight the domain-specificity of the reduced transient distortions measured in Dyslexic participants. Our results proved that saccadic compression of space similarly occurred in both Dyslexic and typical-reading children, a result which is further strengthened by the extensive literature testifying impaired temporal processing as a sign of Dyslexia’s clinical framework.

To summarize our conclusions, with the current study, we brought evidence that dyslexic children failed to modulate visual information shortly before a saccade if temporal processing is involved. Notably, we found that oscillations in temporal sensitivity within the delta band preceded the saccade’s onset in typical-reading children, while a similar fluctuation in perception was absent in dyslexic participants. Such a lack of oscillations can account for the mismatch between the start of the eye movement and the anticipatory shift of the receptive fields, which are primarily involved in determining saccadic transient distortions in space and time. For the first time, we are thus bringing evidence that dyslexic individuals might be inefficient at anticipating saccadic movements at the physiological level, which can represent a concurrent cause in determining any reading impairment.

There is no doubt that Dyslexia is an overly complex clinical condition. Since the last century, its investigation originated a plethora of intriguing and elegant interpretations, spanning from the acquisition of phonological awareness to the magnocellular theory of Dyslexia. Now, our results suggest that it might be time to focus on the receptive field’s predictive remapping as well.

## Materials and Methods

### Participants

A total of 51 children (Age: 127.14 ± 20.27 months, 22 females) participated in this study. 26 of them were diagnosed with Dyslexia (Age: 127.57 ± 21.67 months, 13 females), while the rest were included in the typical-reading group (Age: 125.68 ± 19.85 months, 9 females). Dyslexic children were recruited via a local rehabilitation center which also provided the legal, accredited diagnosis certification according to Italian law. Within the current study, we selected only dyslexic children who were still not included in a rehabilitation program. Typical-reading participants were recruited by contacting the parents using the Istituto Italiano di Tecnologia mailing list. All participants and their parents gave informed consent before participation in the study, which was performed in accordance with the local ethical committee (Comitato Etico Regione Liguria, Genoa, Italy; Prot. IIT_UVIP_COMP_2019 N. 02/2020, 4 July 2020) and the declaration of Helsinki. Demographic information, including reading speeds, are reported in Table T1. Reading speeds were evaluated using the MT-2 reading test^58^, which is one of the most widespread used to support the diagnosis of Dyslexia in Italy, and are reported in syllables per second.

**Table T1.**
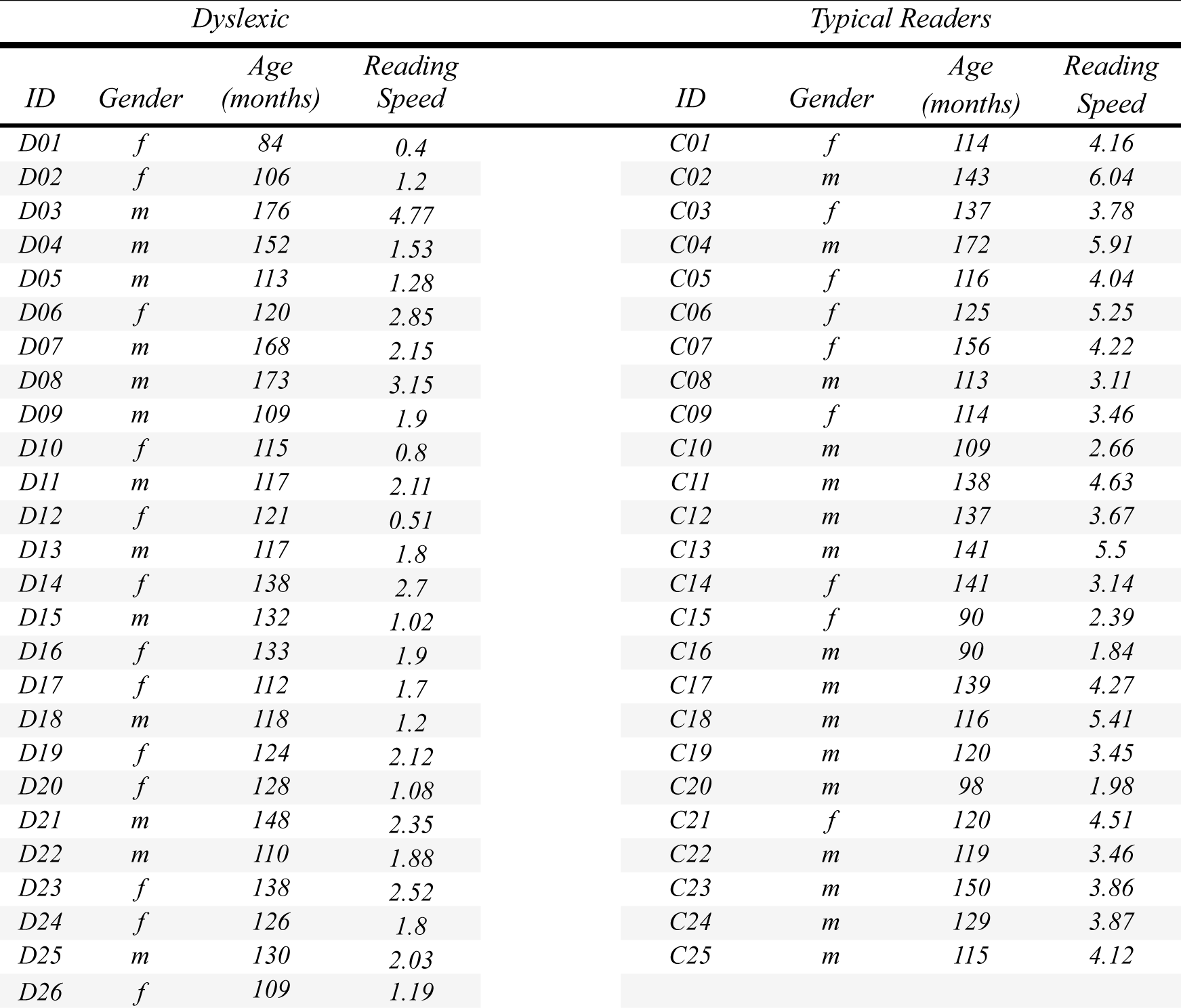
Demographic information for both groups involved in the study.

| *Dyslexic* |  |  |  | *Typical Readers* |  |  |  |
| --- | --- | --- | --- | --- | --- | --- | --- |
| *ID* | *Gender* | *Age (months)* | *Reading Speed* | *ID* | *Gender* | *Age (months)* | *Reading Speed* |
| D01 | f | 84 | 0.4 | C01 | f | 114 | 4.16 |
| D02 | f | 106 | 1.2 | C02 | m | 143 | 6.04 |
| D03 | m | 176 | 4.77 | C03 | f | 137 | 3.78 |
| D04 | m | 152 | 1.53 | C04 | m | 172 | 5.91 |
| D05 | m | 113 | 1.28 | C05 | f | 116 | 4.04 |
| D06 | f | 120 | 2.85 | C06 | f | 125 | 5.25 |
| D07 | m | 168 | 2.15 | C07 | f | 156 | 4.22 |
| D08 | m | 173 | 3.15 | C08 | m | 113 | 3.11 |
| D09 | m | 109 | 1.9 | C09 | f | 114 | 3.46 |
| D10 | f | 115 | 0.8 | C10 | m | 109 | 2.66 |
| D11 | m | 117 | 2.11 | C11 | m | 138 | 4.63 |
| D12 | f | 121 | 0.51 | C12 | m | 137 | 3.67 |
| D13 | m | 117 | 1.8 | C13 | m | 141 | 5.5 |
| D14 | f | 138 | 2.7 | C14 | f | 141 | 3.14 |
| D15 | m | 132 | 1.02 | C15 | f | 90 | 2.39 |
| D16 | f | 133 | 1.9 | C16 | m | 90 | 1.84 |
| D17 | f | 112 | 1.7 | C17 | m | 139 | 4.27 |
| D18 | m | 118 | 1.2 | C18 | m | 116 | 5.41 |
| D19 | f | 124 | 2.12 | C19 | m | 120 | 3.45 |
| D20 | f | 128 | 1.08 | C20 | m | 98 | 1.98 |
| D21 | m | 148 | 2.35 | C21 | f | 120 | 4.51 |
| D22 | m | 110 | 1.88 | C22 | m | 119 | 3.46 |
| D23 | f | 138 | 2.52 | C23 | m | 150 | 3.86 |
| D24 | f | 126 | 1.8 | C24 | m | 129 | 3.87 |
| D25 | m | 130 | 2.03 | C25 | m | 115 | 4.12 |
| D26 | f | 109 | 1.19 |  |  |  |  |

All participants had normal or corrected-to-normal vision, and no child was born prematurely. Data were collected at the rehabilitation center and the Istituto Italiano di Tecnologia in a dimly lit room used for data collection purposes. Although the two groups were balanced, prioritizing age, the age itself was always included in all statistical analyses as a covariate to control the possibility that any effect found was due to typical ongoing developmental processes.

### Apparatus

Stimuli were generated via Matlab v. 2020b through the Psychtoolbox routines and were displayed on a 24” Asus VG248QE LCD monitor with a resolution of 1920 x 1080 pixels and a refresh rate of 144 Hz. Participants sat in front of the screen at a viewing distance of 80 cm, with their heads placed on a chinrest to ensure stable monitoring of eye movements. Saccades were recorded with a Tobii ProSpectrum external eye-tracker (https://www.tobii.com/products/eye-trackers/screen-based/tobii-pro-spectrum), with a sampling rate of 1200 Hz.

### Procedure

To ensure the young participants’ involvement, we created a gamified version of a series of experimental designs already used to investigate saccadic compressions in adults^10,12^. Saccades were always performed from left (−10° of visual angle considering the participants’ head position) to right (+10° of visual angle considering the participants’ head position) and, in both the familiarization and the experimental task, stimuli were green, near equiluminant bars displayed on a red background.

### Familiarization

At the beginning of the experimental session, all participants were introduced to a familiarization procedure consisting of 20 trials. Familiarization was implemented so that participants could acquire confidence with the setup, the experimenter, and the eye movements to perform. In each familiarization trial, a first fixation point appeared at 10° of visual angle to the left of the participant’s head position. After participants had maintained fixation for one second, the point disappeared, and a saccadic target was displayed, 20° of visual angle rightward on the same vertical coordinate. Participants were instructed to move their eyes to the target as quickly as possible. During the familiarization phase, we also collected the average saccadic latency of each participant so that, in the other experimental conditions, we could use them to time the presentation of the stimuli around the saccade’s onset as precisely as possible.

### Temporal Task

In the temporal task (Figure 1), an empty test interval delimited by two sequentially displayed green, vertical bars (subtending 2° x 10° of visual angle) was presented around the fixation point, with the two bars being displayed 3° of visual angle to its right and left. Test intervals were displayed after participants had maintained fixation on the fixation point for at least one second. Vertical bars were separated by a temporal interval ranging from 80ms to 1000ms and were displayed for one single frame (∼7ms) either before (∼-300ms, pre-saccadic condition) or around (∼0ms, peri-saccadic condition) the saccade’s onset. The test interval was defined through two adaptive QUEST^59^ algorithms (range=0.2, tGuessSd=1.2, pThreshold=0.5, β=3.5, ε=0.01, γ=0), one for each trial type (pre-saccadic vs. peri-saccadic). After the execution of the saccadic movement and participant had having maintained fixation on the saccadic target for a random interval between 1000 and 1500ms, a reference interval lasting 230ms was displayed around the saccadic target. In order to conclude the trial, participants were asked to report which interval lasted longer by pressing one of two buttons on the keyboard (“*a*” = test interval, “*l*” = reference interval).

Each condition type was repeated 30 times for a total of 60 possible trials. While for most children such an amount of trial was sufficient to obtain an adequate number of both apo- and perisaccadic trials, for some others we needed to collect additional trials. This was due to the fact that children sometimes failed to perform any trial around the saccadic onset (see *Trial Quantification and Statistical Analysis*).

### Spatial Task

In the spatial task (Figure 4), a green vertical bar (subtending 2° x 34° of visual angle) was displayed for one frame (∼7ms) after a random interval – either approximately before, during, or after the saccade’s onset (−300ms, 0ms, +300ms in reference to the saccade’s onset, respectively). The bar was placed in one of three possible positions (although participants were unaware of this), considering the observer as a reference: −10°, 0°, or +15° degrees of visual angle (with negative values indicating that stimuli appeared leftward to the participant). After successfully performing the saccadic movement, the saccadic target disappeared, and a static green vertical bar appeared on the screen. Participants were instructed to move the bar with the mouse and estimate the position of the first bar. The subsequent trial started once the participants confirmed the response through a mouse click.

Each possible combination of spatial position and temporal displacement was repeated eight times for a total of 72 possible trials.

### Trial quantification and statistical analysis

Before investigating the effect of saccades on temporal perception, we analyzed individual performances scanning for both valid and invalid trials. Specifically, we excluded trials in which it was impossible to quantify the temporal distance between the saccade’s onset and the presentation of the stimuli (e.g., due to missing crucial gaze data).

For the temporal task, within the dyslexic group, we obtained on average 57 ± 9 valid trials while within the typical-reading group we obtained, on average, 58 ± 6 trials. Two typical-reading participants were excluded from the aggregate analysis on the temporal task because fitting psychometric functions into their individual data resulted in negative JNDs (i.e., the psychometric curve was reversed), suggesting that they were not able to understand the task’s assignment (see *Participants’ pruning in the temporal task* section).

It is relevant to note that, during the spatial task, in some trials (∼15% of the perisaccadic trials) participants verbally reported that they were not able to perceive any stimulus. Interestingly, this did not occur for the temporal task, in which – at least, according to what reported by all children – they always perceived both the test and the reference intervals. To mark such trials, participants were told to place the bar controlled via the cursor on the left border of the monitor, so that the response was set to 0. For this reason, all responses included within 0 (left border of the monitor) and five degrees of visual angle were discarded. No difference between groups was found when considering how many times such kind of response was provided (t(49)=-1.17, p=0.25).

We argue that this result might indicate saccadic magnocellular suppression occurring during the task, which should be controlled for using equiluminant stimuli and the red color as a background. However, not only there is some evidence suggesting that magnocellular cells activate in response to a red color^60^, but also achieving equiluminance on a whole stimulus is virtually impossible, at least for two reasons: first, perceived luminance depends on the responses of various retinal cells, whose ratio naturally chances along the retina. Second, chromatic aberrations often arise at the border of a stimulus, thus determining luminance borders that nullify the physical equiluminance on the monitor^61^. In light of these considerations, after excluding the appropriate number of trials the influence of the magnocellular saccadic suppression was minimized and controlled.

In the end, we obtained on average 56 ± 9 trials in the dyslexic group, while we obtained, on average, 48 ± 12 trials in the typical-reading group.

### Fitting Procedures

To evaluate temporal performance, we fitted a cumulative Gaussian function into aggregated bin’s data (Figure 3) following the formula:

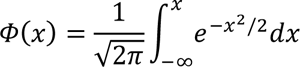

Then, in order to find the best achievable fit, we referred to the maximum likelihood method as described by Watson^62^ and performed 500 bootstrap repetitions per fit. As an index of temporal accuracy, we considered the curve’s inflection point – that is the point of the curve denoting chance level performance with two possible alternatives, better known as the Point of Subjective Equality (PSE). The PSE was also a good indicator of whether participants experienced a compression of perceived time, as higher PSE values corresponds to higher test durations that are matched with the fixed duration of the reference. For instance, a PSE of 350ms in our study indicates a significant compression of perceived time, so that it takes a test lasting 350ms to match the 230ms duration of the reference interval. Due to a technical problem that occurred when performing the experiment with some of the participants (see Appendix 2, “*Fluctuations of the Reference’s Duration*”), as well as for readability purposes, in Figure 3 we directly reported the results of the fitting procedure expressed as a function of the temporal difference between the two intervals.

To evaluate oscillations in performance, we first binned aggregated data into different timestamps, covering up to 1000ms before the saccade’s onset and centred at every −50ms (−50, −150, −250, −350, −450, −550, −650, −750, −850, and −950ms in respect to the saccade’s onset).

We then fitted the best sine function into the binned dataset, using the canonical formula:

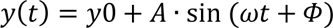

Where *y0* is the vertical shift, *A* is the amplitude, *ω* is the angular frequency (corresponding to *2πf*, with *f* being the ordinary frequency), and Φ is the phase shift. Best-fitting parameters for *y0*, *A*, *ω* and *Φ* were obtained via least-squares estimation method.

The best fitting parameters to approximate typical-reading participants’ performance were obtained through nonlinear least-squares regression (using the Matlab *nlinfit* function) and were the following: *y0* = 0.7535*, A* = 0.0526*, ω* = 431.6282, Φ = 272.3201. Overall, the resultant fit was significantly better than the function y = constant (*R*^2^=0.917, p=0.0012). The best fitting parameters found considering dyslexic participants’ performance, reported only for illustrative purposes as the fit did not converge, were the following: *y0* = - 1851.1281*, A* = −1851.8654*, ω* = −1443.253, Φ = *-*2959.085.

After the best function approximating our empirical data was defined, we proceeded to test the reliability of our predictions by comparing its empirical *R*^2^ with an ensemble of comparable *R*^2^ obtained through a distribution of surrogate datasets. Surrogate datasets were generated via random sampling, starting from the original observations. To generate surrogate data, from the original matrix we randomly mixed the ‘time from saccade’ index for each trial, so to maintain similar time distributions and avoid spurious effects linked to different time stamps. Then, we replicated the analysis performed on the original dataset, and obtained a *R*^2^ measure indicating the correspondence between that specific iteration of the surrogate data and the predicted values obtained through the best sine. To correct for multiple comparisons, for each sine function fitted on the surrogated dataset *y0*, *A*, ω, *Φ* were treated as free-to-vary parameters. After 10,000 bootstrapping repetitions, we compared the *R*^2^ measured using the empirical data against the *R*^2^ distribution obtained with the 10,000 surrogate datasets. Lastly, to prove that the empirical *R*^2^ was not stochastically determined, we measured the proportion of times that a higher *R*^2^ was found in the surrogate distribution as follows:

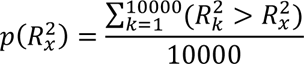

If 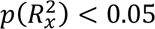, we then concluded that the R^2^ obtained with the empirical data was statistically different (in our case, higher) from an *R*^2^ using the same dataset.

All fitting procedures were performed using Matlab 2020b (https://mathworks.com), while statistical analyses were performed using JASP v0.17.2.1 (https://jasp-stats.org/).

### Participants’ pruning in the temporal task

Two participants in the typical reading group were excluded from the aggregate analysis, namely C15 and C17, due to their inability to properly complete the task. As showed in Figure 7, the psychometric curve for both participants resulted reversed, meaning that the task was not properly understood or executed. No other participant showed similar behavior.

**Figure 7:**
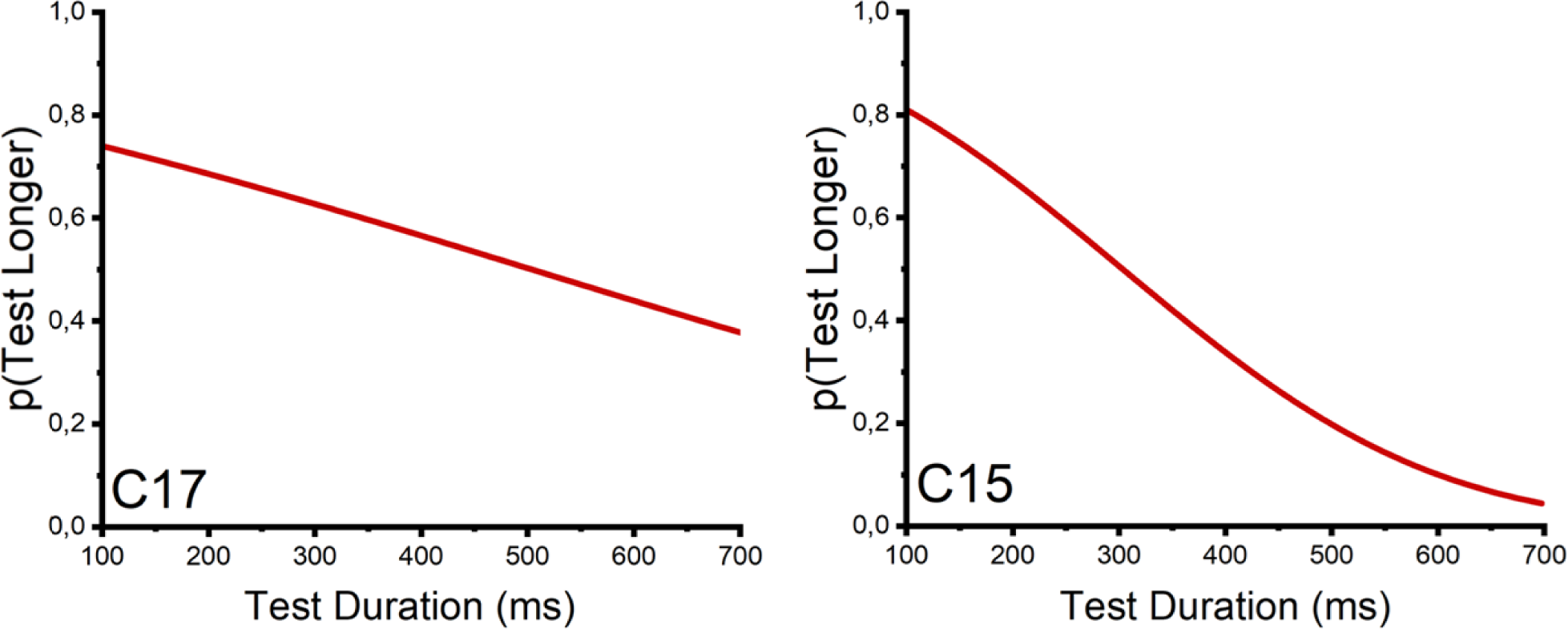
Psychometric curves of the two participants excluded from the aggregated analysis. In both cases, the curve is reversed– suggesting that participants did not understand or executed the task.

### Posterior odds for the logistic regression

In the Result section, we reported the results of the logistic regression performed on the correctness of responses, considering as a covariate the Age and the Reading Speed, and as factors both the Group (Typical-reading vs. Dyslexic) and bin (−950 vs. −850 vs. −750 vs. −650 vs. −550 vs. −450 vs. −350 vs. −250 vs. −150 vs. - 50ms vs. 50ms). Here we report in Table T2 all the relevant parameters for all tested models, while in Table T3 we reported the posterior summary for all coefficients.

**Table T2:** Model Comparison against the null model. The table reports the prior model probabilities P(M), the posterior model probabilities P(M|data), the posterior model odds BFM, and the Bayes Factor BF10.

| Models | $P(M)$ | $P(M\vert data)$ | $BF_M$ | $BF_{10}$ |
| --- | --- | --- | --- | --- |
| Null model (incl. Age and Reading Speed) | 0.458 | 0.128 | 0.173 | 1.000 |
| bin+Group | 0.458 | 0.789 | 4.427 | 6.183 |
| Bin | 0.042 | 0.075 | 1.861 | 6.451 |
| Group | 0.042 | 0.008 | 0.19 | 0.708 |

**Table T3:**
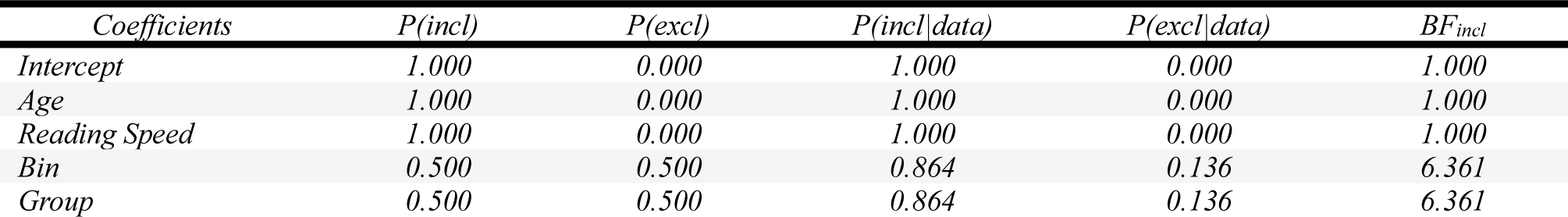
Posterior Summaries of coefficients. The table reports the prior inclusion probability P(incl), the prior exclusion probability P(excl), the posterior inclusion probability(incl|data), the posterior exclusion probability P(excl|data), and the change from prior inclusion odds to posterior inclusion odds BFincl for each component, averaged by all the models that includes that component.

| Coefficients | $P(incl)$ | $P(excl)$ | $P(incl\vert data)$ | $P(excl\vert data)$ | $BF_{incl}$ |
| --- | --- | --- | --- | --- | --- |
| Intercept | 1.000 | 0.000 | 1.000 | 0.000 | 1.000 |
| Age | 1.000 | 0.000 | 1.000 | 0.000 | 1.000 |
| Reading Speed | 1.000 | 0.000 | 1.000 | 0.000 | 1.000 |
| Bin | 0.500 | 0.500 | 0.864 | 0.136 | 6.361 |
| Group | 0.500 | 0.500 | 0.864 | 0.136 | 6.361 |

**Table T4:**
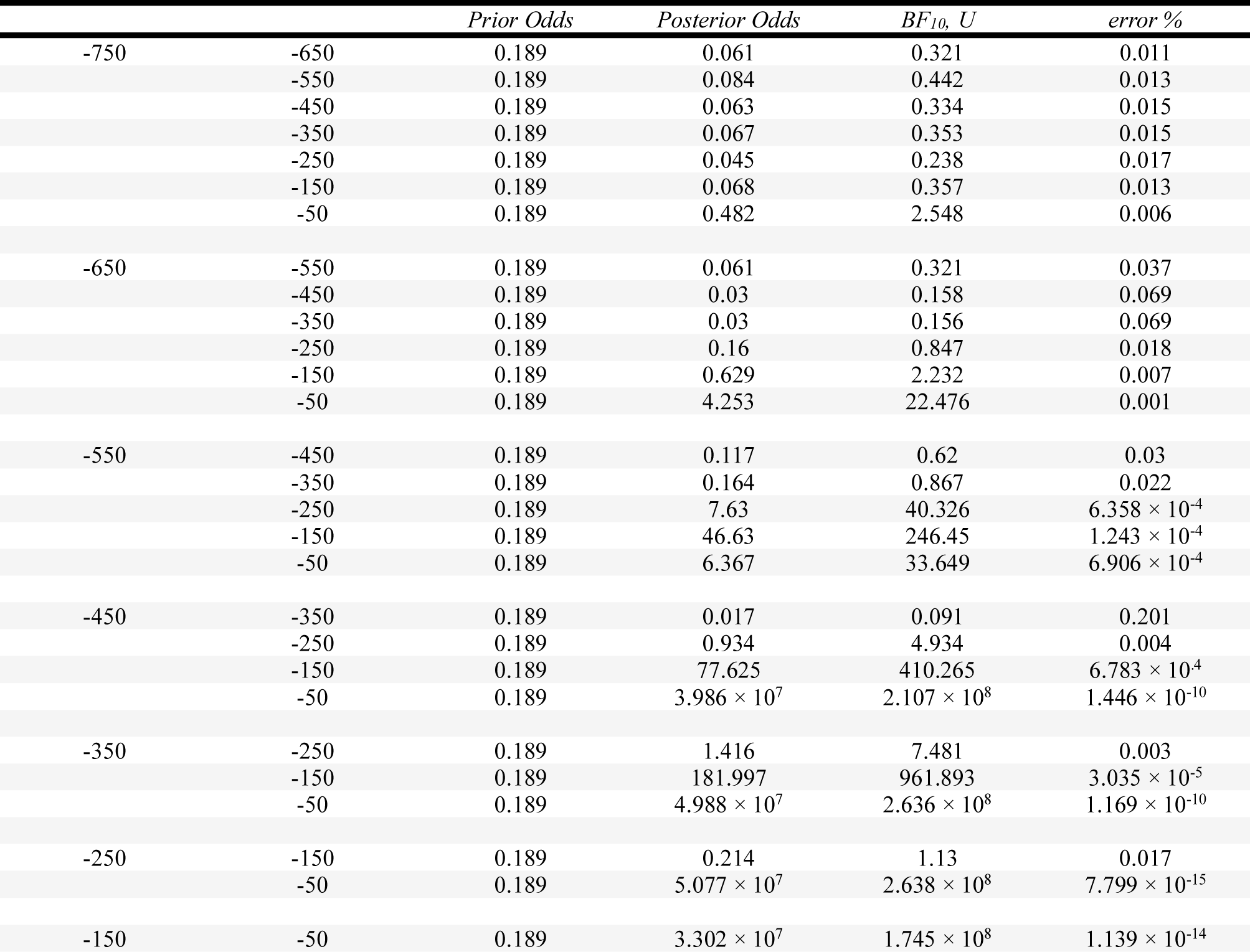
Post-hoc analyses following the Bayesian ANCOVA performed over the factor Bin. The posterior odds have been corrected for multiple comparisons by fixing to 0.5 the prior probability that the null hypothesis holds across all tests. “BF10, U” indicates the uncorrected Bayes Factor, while “error %” indicates the error of the Gaussian quadrature integration routine used for the evaluation of the BF. Note that calculation errors below 10% do not hinder the interpretation of the comparison (within our analysis, this always holds true but for the comparison between −450 and −350ms).

|  |  | *Prior Odds* | *Posterior Odds* | *BF<sub>10</sub>, U* | *error %* |
| --- | --- | --- | --- | --- | --- |
| -750 | -650 | 0.189 | 0.061 | 0.321 | 0.011 |
|  | -550 | 0.189 | 0.084 | 0.442 | 0.013 |
|  | -450 | 0.189 | 0.063 | 0.334 | 0.015 |
|  | -350 | 0.189 | 0.067 | 0.353 | 0.015 |
|  | -250 | 0.189 | 0.045 | 0.238 | 0.017 |
|  | -150 | 0.189 | 0.068 | 0.357 | 0.013 |
|  | -50 | 0.189 | 0.482 | 2.548 | 0.006 |
| -650 | -550 | 0.189 | 0.061 | 0.321 | 0.037 |
|  | -450 | 0.189 | 0.03 | 0.158 | 0.069 |
|  | -350 | 0.189 | 0.03 | 0.156 | 0.069 |
|  | -250 | 0.189 | 0.16 | 0.847 | 0.018 |
|  | -150 | 0.189 | 0.629 | 2.232 | 0.007 |
|  | -50 | 0.189 | 4.253 | 22.476 | 0.001 |
| -550 | -450 | 0.189 | 0.117 | 0.62 | 0.03 |
|  | -350 | 0.189 | 0.164 | 0.867 | 0.022 |
| | -250 | 0.189 | 7.63 | 40.326 | $6.358 \times 10^{-4}$ |
| | -150 | 0.189 | 46.63 | 246.45 | $1.243 \times 10^{-4}$ |
| | -50 | 0.189 | 6.367 | 33.649 | $6.906 \times 10^{-4}$ |
| -450 | -350 | 0.189 | 0.017 | 0.091 | 0.201 |
|  | -250 | 0.189 | 0.934 | 4.934 | 0.004 |
| | -150 | 0.189 | 77.625 | 410.265 | $6.783 \times 10^{-4}$ |
| | -50 | 0.189 | $3.986 \times 10^7$ | $2.107 \times 10^8$ | $1.446 \times 10^{-10}$ |
| -350 | -250 | 0.189 | 1.416 | 7.481 | 0.003 |
| | -150 | 0.189 | 181.997 | 961.893 | $3.035 \times 10^{-5}$ |
| | -50 | 0.189 | $4.988 \times 10^7$ | $2.636 \times 10^8$ | $1.169 \times 10^{-10}$ |
| -250 | -150 | 0.189 | 0.214 | 1.13 | 0.017 |
| | -50 | 0.189 | $5.077 \times 10^7$ | $2.638 \times 10^8$ | $7.799 \times 10^{-15}$ |
| -150 | -50 | 0.189 | $3.302 \times 10^7$ | $1.745 \times 10^8$ | $1.139 \times 10^{-14}$ |

### Post-hoc comparison on spatial mislocalization

## Appendix 1 Eye movement parameters

To define the Time to saccadic onset, in all experiments we calculated the temporal difference between the presentation of the stimulus and the closest eye movement which subtended more than 2° of visual angle. For the temporal task, the reference for the stimulus onset was the second bar that defined the test interval; for the spatial task, the reference for the stimulus onset was naturally the time at which the stimulus appeared.

To expand (and support) our main findings, we decided to investigate potential differences in eye movements between typical-reading and dyslexic children. We thus focused our analysis targeting the saccadic speed measured during both the temporal and the spatial task. For this analysis, we included only trials in which participants performed a saccade which reached 3° prior to the saccadic target (with the latter limit being used also to calculate the speed itself, by simply evaluating the ratio between the gaze distance and time). Due to artifacts characterizing some of the traces, saccadic onsets and offsets were computed on a trial-by-trial basis.

Lastly, to further reduce the influence of artifact within traces, in the analysis we included only speeds that were higher than 200 and lower than 700°/sec.

Comparing saccadic speeds across groups, we found that overall dyslexic children exhibited slower saccades when compared to typical readers, both in the temporal (Dyslexic, M = 406.384 ± 69.436ms; Typical Readers: M = 445.11 ± 64.805ms, BF_10_ = 5.17 × 10^34^) and spatial (Dyslexic, M = 438.416 ± 71.98ms; Typical Readers: M = 449.974 ± 66.694ms, BF_10_ = 34.275) task. Overall, these results align with previous literature showing reduced saccadic speed in Dyslexic children^63,64^.

## Appendix 2 Fluctuations of the Reference’s Duration

Due to a technical issue with the setup, for five participants in the Dyslexic group the duration of the reference interval was centered on 260 rather than 230ms (namely, D02, D05, D06, D10, and D12). To challenge this issue, we evaluated the correctness of each response by directly comparing the physical duration of the test with the one of the reference on a trial-by-trial basis. For similar reasons, in Figure 3 in the main text, we fitted psychometric functions considering the probability of reporting the test as the longest stimulus expressed as a function of the physical difference between the two intervals.

**Appendix 2–Figure 1:**
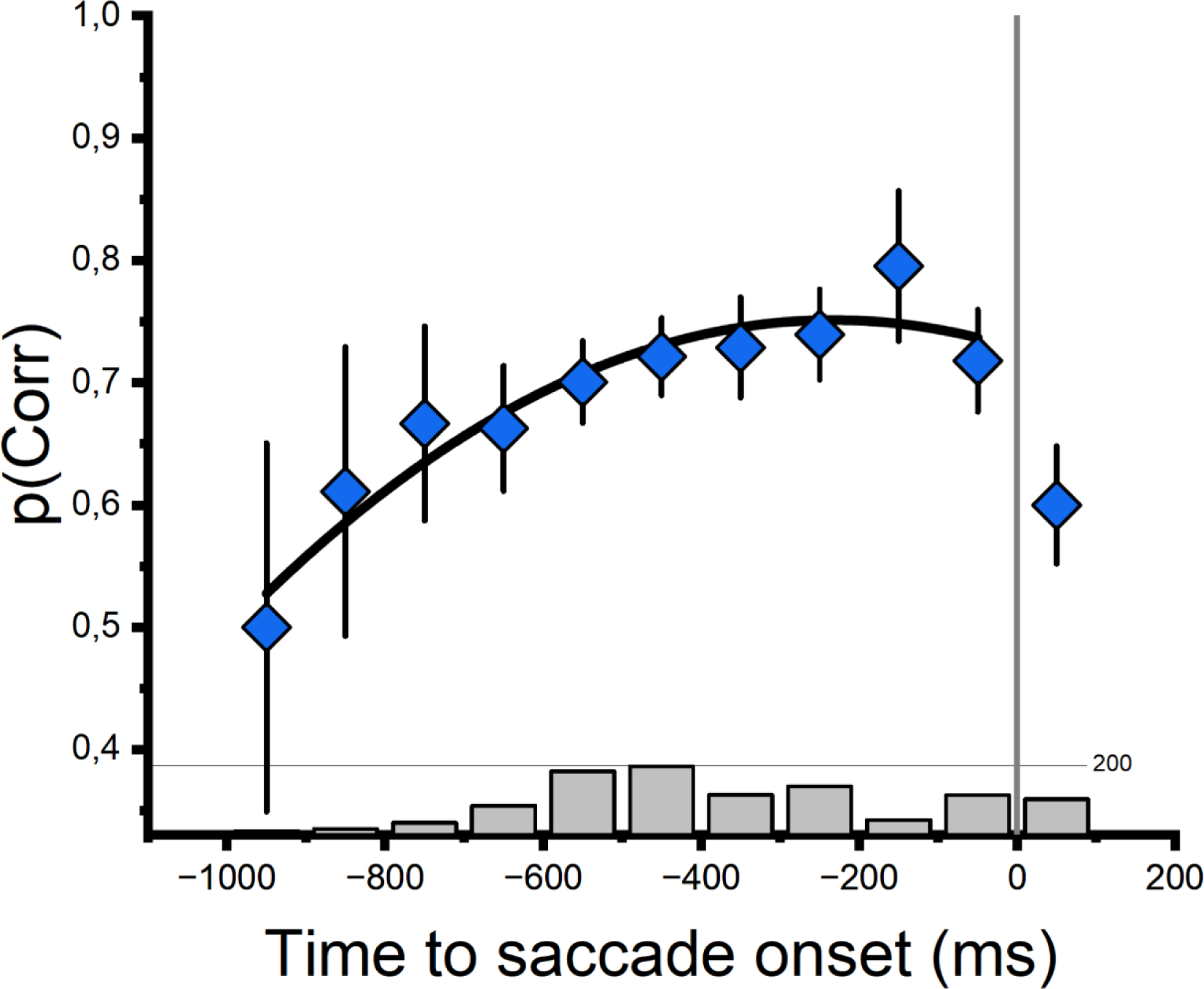
Changes in temporal accuracy prior to the saccadic onset, including Dyslexic participants for whom the reference interval was centered around 230ms (n=21). Error bars indicate ±SEM, while grey bars below each bin indicate the number of trials for that given bin.

Due to this issue, we repeated the analysis excluding the five participants mentioned above (Appendix 2– Figure 1). Even when considering only participants for whom the reference was centered around 230ms, we found that temporal perception before the saccadic onset in Dyslexic children did not show any oscillatory pattern. The nonlinear fit failed to adequately converge, resulting in extremely high parameters (reported here for illustrative purposes only: *y0* =-622.6825, *ω* = −8.41 × 10^4^, Φ = 4.83 × 10^6^, *A* = −623.4365) and confirming that sinusoidal-like changes in timing performances did not anticipate saccades in pre-rehabilitation dyslexic children.

## Acknowledgements

The authors would like to thank David Burr for his fruitful comments during the early stages of the work, as well as Concetta Morrone for her paramount insights provided both before and after having read the first draft of the manuscript. The authors would also like to thank all the children that partook in this study, their families, and all the staff of the Centro Leonardo for their availability and enthusiasm, as without them this study work would not have been possible.

## Data Availability

Data can be accessed at the following link: https://zenodo.org/records/10263273 Gaze data can be accessed at the following link: https://zenodo.org/records/10264504.

